# Phosphoproteomics-based Profiling of Kinase Activities in Cancer Cells

**DOI:** 10.1101/066019

**Authors:** Jakob Wirbel, Pedro Cutillas, Julio Saez-Rodriguez

## Abstract

Cellular signaling, predominantly mediated by phosphorylation through protein kinases, is found to be deregulated in most cancers. Accordingly, protein kinases have been subject to intense investigations in cancer research, to understand their role in oncogenesis and to discover new therapeutic targets. Despite great advances, an understanding of kinase dysfunctioning in cancer is far from complete.

A powerful tool to investigate phosphorylation is mass-spectrometry (MS)-based phosphoproteomics, which enables the identification of thousands of phosphorylated peptides in a single experiment. Since every phosphorylation event results from the activity of a protein kinase, high-coverage phosphoproteomics data should indirectly contain comprehensive information about the activity of protein kinases.

In this chapter, we discuss the use of computational methods to predict kinase activity scores from MS-based phosphoproteomics data. We start with a short explanation of the fundamental features of the phosphoproteomics data acquisition process from the perspective of the computational analysis. Next, we briefly review the existing databases with experimentally verified kinase-substrate relationships and present a set of bioinformatic tools to discover novel kinase targets. We then introduce different methods to infer kinase activities from phosphoproteomics data and these kinase-substrate relationships. We illustrate their application with a detailed protocol of one of the methods, KSEA (Kinase Substrate Enrichment Analysis). This method is implemented in Python within the framework of the open-source Kinase Activity Toolbox (kinact), which is freely available at http://github.com/saezlab/kinact/.

## 1 Introduction

Protein kinases are major effectors of cellular signaling, in the context of which they form a highly complex and tightly regulated network that can sense and integrate a multitude of external stimuli or internal cues. This kinase network exerts control over cellular processes of fundamental importance, such as the decision between proliferation and apoptosis [Jørgensen and Linding, 2010]. Deregulation of kinase signaling can lead to severe diseases and is observed in almost every cancer [Hanahan and Weinberg, 2011]. For instance, a single constitutively active kinase, originating from the fusion of the BCR and ABL genes, can give rise to and sustain chronic myeloid leukemia [Sawyers, 1999]. Accordingly, the small molecule inhibitor of the BCR-ABL kinase, Imatinib, has shown unprecedented therapeutic effectiveness in affected patients [Sawyers et al., 2002].

Fuelled by these promising clinical results, due to the essential role for kinases in the patho-mechanism of cancer, and because kinases are in general pharmacologically tractable [Zhang et al., 2009], a range of new kinase inhibitors has been approved or is in development for different cancer types [Gonzalez de Castro et al., 2012]. However, not all eligible patients respond equally well, and in addition, cancers often develop resistance to initially successful therapies. This calls for a deeper understanding of kinase signaling and how it can be exploited therapeutically [Cutillas, 2015].

By definition, the activity of a kinase is reflected in the occurrence of phosphorylation events catalyzed by this kinase. Thus, analysis of kinase activity was traditionally achieved by monitoring the phosphorylation status of a limited number of sites known to be targeted by the kinase of interest using immunochemical techniques [Bertacchini et al., 2014]. This, however, requires substantial prior-knowledge and yields a comparably low throughput. Other approaches exist, e.g. protein kinase activity assays [Cutillas et al., 2006; Yu et al., 2009] or attempts to measure kinase activity with chromatographic beads functionalized with ATP or small molecule inhibitors [McAllister et al., 2013].

Mass spectrometry-based techniques to measure phosphorylation can identify thousands of phosphopeptides in a single sample with ever increasing coverage, throughput, and quality, nourished by technological advances and dramatically increased performance of MS instruments in recent years [Doll and Burlingame, 2015; Choudhary and Mann, 2010; Sabidó et al., 2012]. High-coverage phosphoproteomics data should indirectly contain information about the activity of many active kinases. The high-content nature of phosphoproteomics data, however, poses challenges for computational analysis. For example, only a small subset of the described phosphorylation sites can be explicitly associated with functional impact [Beltrao et al., 2012].

As a means to extract functional insight, methods to infer kinase activities from phosphoproteomics data based on prior-knowledge about kinase-substrate relationships have been put forward [Qi et al., 2014; Casado et al., 2013; Yang et al., 2015; Mischnik et al., 2015]. The knowledge about kinase-substrate relationships, compiled in databases like Phospho-SitePlus [Hornbeck et al., 2015] or Phospho.ELM [Dinkel et al., 2011], covers only a limited set of interactions. Alternatively, computational resources to predict kinase-substrate relationships based on kinase recognition motifs and contextual information have been used to enrich the collections of substrates per kinase [Horn et al., 2014; Song et al., 2012], but the accuracy of such kinase-substrate relationships has not been validated experimentally for most cases. The inferred kinase activities can in turn be used to reconstruct kinase network circuitry or to predict therapeutically relevant features such as sensitivity to kinase inhibitor drugs [Casado et al., 2013].

In this chapter, we start with a brief description of phosphoproteomics data acquisition, highlighting challenges for the computational analysis that may arise out of the experimental process. Subsequently, we will present different computational methods for the estimation of kinase activities based on phosphoproteomics data, preceded by the kinase-substrate resources these methods use. One of these methods, namely KSEA (Kinase-Substrate Enrichment Analysis), will be explained in more detail in the form of a guided, stepwise protocol, that is available as part of the Python open-source Toolbox kinact (Toolbox for Kinase Activity Scoring) at http://www.github.com/saezlab/kinact/.

## 2 Phosphoproteomics Data Acquisition

For a summary of technical variations or available systems for the experimental setup of phosphoproteomics data acquisition, we would like to refer the interested reader to dedicated publications such as [Riley and Coon, 2016; Nilsson, 2012]. We provide here a short overview about the experimental process to facilitate the understanding of common challenges that may arise for the data analysis that we will focus on.

Mass spectrometry-based detection of peptides with post-translational modifications (PTM) usually requires the same steps, independent of the modification of interest: (i) cell lysis and protein extraction with special focus on PTM preservation, (ii) digestion of proteins with an appropriate protease, (iii) enrichment of peptides bearing the modification of interest, and (iv) analysis of the peptides by LC-MS/MS [Hennrich and Gavin, 2015]. After the experimental work, additional data processing steps are required to identify the position of the modification, e.g. the residue that is phosphorylated. For almost every step, different protocols are available, starting from various proteases for protein digestion to different data acquisition methods for MS [Riley and Coon, 2016].

### 2.1 Phosphopeptide Enrichment

Naturally, the enrichment of phosphopeptides is a pivotal step for phosphoproteomics. Next to the enrichment method used, the choice of the protease [Giansanti et al., 2015] or the MS ionization method [Ruprecht et al., 2016] also seem to have an impact on the part of the phosphoproteome that is sampled. For phosphopeptide enrichment, the field is dominated by immobilized metal affinity chromatography (IMAC) and metal oxide affinity chromatography (MOAC), which all exploit the affinity of the phosphorylation towards metal ions. Popular techniques include Fe3+-IMAC, Ti4+-IMAC [Zhou et al., 2013], or TiO2-MOAC. Alternatively, more traditional biochemical methods involving immunoaffinity purification are also in use for enrichment of phosphopeptides, although these are generally limited to studies of phosphotyrosine [Rush et al., 2005].

Of note, the different enrichment methods show limited overlap in the detected phosphopeptides, although this can also be observed for replicates of runs using the identical enrichment method, as discussed below [Ruprecht et al., 2015].

After enrichment, the phosphopeptides are separated chromatographically, usually by reversed phase liquid chromatography (RPLC), and then enter the mass spectrometer for tandem MS analysis (MS/MS), completing the workflow of LC-MS/MS. Variations in the chromatography method used as well as the multitude of mass spectrometry instrument types are reviewed in detail elsewhere [Riley and Coon, 2016]. Generally, the quality of the chromatographic separation will have a big impact on the number of phosphopeptides that can confidently be identified. Chromatography runs of higher quality also take more time, so that a trade-off between resolution and throughput must be devised for each experiment.

### 2.2 Data Acquisition

For most phosphoproteomics studies so far, the MS instrument is operated in the data-dependent acquisition (DDA) mode. Therein, precursor ions from a first survey scan are selected-typically based on relative ion abundance- in order to generate fragmentation spectra in a second MS run [Domon and Aebersold, 2006], for which a database search yields the corresponding peptide sequences [Nesvizhskii, 2007]. As a result, peptide detection in DDA is on the one hand biased towards high abundance species, but also considerably irreproducible due to stochastic precursor ion selection [Liu et al., 2004]. This inherent under-sampling of DDA usually leads to missing data points in LC-MS/MS datasets. However, this problem may be solved to some extent by extracting ion chromatograms of the peptides that are missing in some of the runs that are being compared [Cutillas and Vanhaesebroeck, 2007; Cutillas et al., 2005; Bateman et al., 2014; Alcolea et al., 2012], by matching across samples [Cox et al., 2014], or with the accurate mass and retention tag method [Strittmatter et al., 2003].

In an alternative operation mode, selected reaction monitoring/multiple reaction monitoring (SRM/MRM), the presence and abundance of only a limited set of pre-specified peptides with known fragmentation spectra is surveyed [Lange et al., 2008]. This targeted approach overcomes many of the issues of shotgun methods, but is usually not feasible for large-scale investigation of the complete phosphoproteome.

Data-independent acquisition (DIA), e.g. SWATH-MS [Gillet et al., 2012] tries to address the shortcoming of both established data acquisition strategies in order to combine the throughput of DDA with the reproducibility of SRM. In DIA, fragmentation spectra are generated for all precursor ions in a specific window of *m/z* ratios, leading to a complete map of fragmentation spectra, followed by computational extraction of quantitative information for known spectra. For phosphoproteomics, DIA-MS has already been applied to investigate insulin signaling [Parker et al., 2015] or histone modifications [Sidoli et al., 2016]. However, the spectra generated by DIA-MS are usually highly complex and require intricate data extraction techniques, which is even more challenging for modified peptides. Recently, a computational resource for the detection of modified peptides has been put forward [Keller et al., 2016]. Overall, the available methods for DIA have as yet to mature in order to challenge the use of DDA in large-scale studies of the phosphoproteome [Riley and Coon, 2016].

### 2.3 Quantitative Phosphoproteomics

As for regular proteomics, several experimental methods or post-acquisition tools exist to quantitate detected phosphopeptides. Those can roughly be divided into isotope labeling and label-free quantitation. In general, stable isotope labeling requires more experimental effort than label-free quantitation, but at the same time enables multiplexing of samples with different isotopes or combinations.

Stable isotope labeling by metabolic incorporation of amino acids (SILAC) is mainly used for cell cultures, in the medium of which different stable isotopes are provided that will be incorporated into the proteins of the cells. At the point of analysis, cell extracts are mixed and then jointly investigated with mass spectrometry. Mass differences between peptide pairs due to the isotopic labeling can be exploited for relative quantitation [Ong et al., 2002]. Currently, up to three conditions (light, medium, heavy) can be multiplexed. Further developments of SILAC even produced an in-vivo SILAC mouse model for the proteomic and phosphoproteomic analysis of skin cancerogenesis [Zanivan et al., 2013] and super-SILAC for the analysis of tissues [Shenoy and Geiger, 2015], in which a metabolically labeled, tissue-specific protein mix from several cell lines, representing the complexity of the investigated proteome, is mixed with the tissue lysate as internal standard for quantification.

Chemical modification of peptides with tandem mass tags (TMT) or isobaric tags for relative and absolute quantitation (iTRAQ) are two different methods based on tags with reactive groups that bind to peptidyl amines in the peptides after protein digestion. Again, different samples are mixed before mass spectrometry analysis, whereas for TMT or iTRAQ up to 8 samples can be multiplexed. In the first MS run, the peptides with different isobaric tags are indistinguishable, but upon fragmentation in the second MS run, each tag generates a unique reporter ion fragmentation spectrum, which can be used for relative quantitation of the tagged peptides [Thompson et al., 2003; Ross et al., 2004].

Label-free quantitation (LFQ), on the other hand, relies mainly on post-acquisition data analysis, so that no modification of the essential experimental workflow needs to be implemented. Comparison of an -in theory-unlimited number of different samples is therefore possible, which is associated with the downside of prolonged analysis time as multiplexing samples is not possible. While label-free approaches usually provide a deeper coverage of the proteome than label-based methods, the reproducibility and precision of quantification are inferior, so that more technical replicates are needed for confident quantification in LFQ [Li et al., 2012]. Typically, label-free quantitation is achieved by integration of peak area measurements, i.e. the area under the curve, for individual peptides [Chelius and Bondarenko, 2002] or spectral counting, which reflects that the probability to sample more abundant peptides is higher [Neilson et al., 2011].

For the case of phosphoproteomics, in contrast to regular proteomics, an additional challenge for quantitation arises from the fact that information from different peptides of the same protein cannot be integrated. While in regular proteomics the abundances of every peptide in the protein can be combined, the quantitation of a single phosphosite depends on direct measurements of peptides with the specific modification. Therefore, the sample sizes in phosphoproteomics quantitation are much smaller and can even consist of the measurement of only a single peptide.

Furthermore, different phosphosites within the same protein will most probably not show similar pattern of phosphorylation. This may give rise to problems for subsequent analysis, if this analysis is conducted on protein rather than on phosphosite level.

### 2.4 Phosphosite Assignment

Phosphopeptides in large-scale phosphoproteomics experiments are identified from LC-MS/MS runs by interpreting MS/MS spectra using a suitable search engine. Several of such search engines now exist; popular ones include Mascot, Sequest, Protein Prospector and Andromeda [Perkins et al., 1999; Clauser et al., 1999; MacCoss et al., 2002; Cox et al., 2011]. The process of determining peptide sequences from MS/MS data involves matching the mass to charge ratios of fragment ions in MS/MS spectra to the theoretical fragmentation of all protein-derived peptides in protein databases. Depending on the organism being investigated, protein databases from UniProt or NCBI are used. Each search engine has its own scoring system to reflect the confidence of peptide identification, which is a function of MS and MS/MS spectral quality. False discovery rate (FDR) may be determined by performing parallel searches against scrambled or reversed protein databases containing the same number of sequences as the authentic protein database. The FDR is then calculated as the ratio of positive peptide identifications in the decoy database divided by those derived from the forward search. A FDR of 1% at the peptide level is normally considered adequate.

Deriving peptide sequences with these methods is a relatively straightforward process. However, site localization can be a problem when peptide sequences contain more than one amino acid residue that can be phosphorylated. To address this problem, several methods to determine precise localization of phosphorylation within a phosphopeptide have been published. Ascore uses a probabilistic approach to assess correct site assignment [Beausoleil et al., 2006] and the algorithm has been applied alongside the Sequest search engine. The Mascot delta score, introduced by the Kuster group, simply determines the differences in Mascot scores between the different possibilities for phosphosite localization within a phosphopeptide [Savitski et al., 2011]. The larger the delta score, the greater the probability of correct site assignment. Other similar methods have been published [Chalkley and Clauser, 2012] and some of them are now incorporated into search engines [Baker et al., 2011]. The output of the phosphopeptide identification step generally contains scores for both probability of correct peptide sequence identification as well as phosphosite localization.

### 2.5 Pifalls in the Analysis of MS-based Phosphoproteomics Data

Although the available experimental methods for MS-based phosphoproteomics data acquisition have evolved considerably in the last years, leading to a steadily increasing number of detected phosphosites, several limitations remain for the investigation of signaling processes using phosphoproteomics data.

While it has been estimated that there are around 500,000 phosphorylation sites in the human proteome [Lemeer and Heck, 2009], the number of phosphosites that can be identified in a single MS experiment usually ranks around 10,000 to up to 40,000 [Sharma et al., 2014]. Therefore, the sampled phosphoproteomic picture is incomplete. It has to be taken into account though, that not all possible phosphorylation sites are expected to be modified at the same time point, because their regulation is context-dependent, for example, some are controlled differentially at each stage of the cell cycle, while others only change under specific external stimulation mediated by exposition to growth factors or other effector molecules. The hope is therefore, that a significantly larger portion of phosphosites could be achieved with improving technology and by increasing the diversity of biologically relevant conditions analyzed to study the role of phosphoregulation in cell signaling. So far though, in different MS runs or replicates, usually a distinct set of phosphosites is detected, as the selection of precursor ions is stochastic. This leads to incomplete datasets with a high number of missing data points, challenging computational investigation of the data like clustering or correlation analysis. However, as discussed above, approaches in which phosphopeptide intensities are compared across MS runs post-acquisition minimizes this problem [Alcolea et al., 2012].

The functional impact of a phosphorylation event is known only in the minority of cases [Beltrao et al., 2012]. Indeed, it has been hypothesized that a substantial fraction of phosphorylation sites are non-functional [Landry et al., 2009], since phosphorylation sites tend to be poorly conserved throughout species [Beltrao et al., 2009]. Although approaches to study the function of individual phosphorylation events have been proposed [Beltrao et al., 2013], it may be that a large part of the detected phosphosites serves no function at all. Thus, non-functional sites add a substantial amount of noise to phosphoproteomics data and complicate the computational analysis.

The inference of kinase activity from phosphoproteomics data that will be described in the next section aims to overcome these limitations, by integration of the information from many phosphosites, along prior knowledge on kinases-substrate relationships, into a single measure for the kinase activity. However, a general caveat determined by the experimental workflow remains for the inference of kinase activities. Since highly abundant precursor ions are more likely to be selected for fragmentation and therefore detection, all methods preferentially detect highly active kinases.

## 3 Computational Methods for Inference of Kinase Activity

Traditionally, biochemical methods have been common to study kinase activities *in vitro* and are still broadly used [Newman et al., 2014; Glickman, 2012]. However, on the one hand those methods are generally limited in throughput and time-consuming. On the other hand *in vitro* methods might not accurately reflect the *in vivo* activities of kinases in a specific cellular context. MS-based methods have also been applied for assays of kinase activity [Cutillas et al., 2006; Yu et al., 2009], in which the abundance of known target phosphosites are monitored by MS after an *in vitro* enzymatic reaction.

Since every phosphorylation event results –by definition- from the activity of a kinase, phosphoproteomics data should be suitable to infer the activity of many kinases from comparably low experimental effort. This task requires computational analysis of the detected phosphorylation sites (phosphosites), since thousands of phosphosites can routinely be measured in a single experiment. Several methods have been proposed in recent years, all of which utilize prior-knowledge about kinase-substrate interactions, either from curated databases or from information about kinase recognition motifs.

### 3.1 Resources for Kinase-Substrate Relationships

As the large-scale detection of phosphorylation events using mass spectrometry became routine, many freely available databases that collect experimentally verified phosphosites have been set up, including PhosphoSitePlus [Hornbeck et al., 2015], Phospho.ELM [Dinkel et al., 2011], Signor [Perfetto et al., 2016], or PHOSIDA [Gnad et al., 2011], to name just a few. The databases differ in size and aim; PHOSIDA for example provides a tool for prediction of putative phosphorylation or recently also acetylation sites. Phospho.ELM computes a score for the conservation of a phosphosite and Signor is focused on interactions between proteins participating in signal transduction. The arguably most prominent database for expert-edited and curated interactions between kinases and individual phosphosites (that have not been derived from *in vitro* studies) is PhosphoSitePlus, currently encompassing 16,486 individual kinase-substrate relationships [04-2015]. For *Saccharomyces cerevisiae*, the database PhosphoGRID provides analogous information [Sadowski et al., 2013]. PhosphoNetworks also contains information about kinase-substrate interactions, but on protein, not on single phosphosite level [Hu et al., 2014]. Specific information about targets of phosphatases can be found in DEPOD [Duan et al., 2015]. Also in the Phospho.ELM database, phosphosites have been associated with regulating kinases, although this information is available for only about 10% of the 37,145 human phosphosites in the database [04-2015].

As it has been estimated that there are between 100,000 [Zhang et al., 2002] and 500,000 [Lemeer and Heck, 2009] possible phosphosites in the human proteome, the evident low coverage of the curated databases motivated the development of computational tools to predict *in vivo* kinase-substrate relationships. These methods identify putative new kinase-substrate relationships based on experimentally derived kinase recognition motifs, which was pioneered by Scansite [Obenauer et al., 2003] that uses position-specific scoring matrices (PSSMs) obtained by positional scanning of peptide libraries [Chen and Turk, 2010] or phage display methods [Sidhu and Koide, 2007]. Another approach, Netphorest [Miller et al., 2008] tries to classify phosphorylation sites according to the relevant kinase family instead of predicting individual kinase-substrate links. However, the *in vitro* specificity of kinases differs significantly from the kinase activity inside of the cell, biasing the experimentally identified kinase recognition motifs [Hjerrild et al., 2004]. The integration of contextual information, for example co-expression, protein-protein interactions, or subcellular co-localization, markedly improves the accuracy of the predictions [Newman et al., 2014]. The software packages NetworKIN [Linding et al., 2008] (recently extended in the context of the resource KinomeXplorer [Horn et al., 2014], correcting for biases caused by over-studied proteins) and iGPS [Song et al., 2012] are examples for methods that combine information about kinase recognition motifs, *in vivo* phosphorylation sites, and contextual information, e.g. from the STRING database [Szklarczyk et al., 2015]. Recently, Wagih et al. presented a method to predict kinase specificity for kinases without any known phosphorylation sites [Wagih et al., 2016]. Based on the assumption that functional interaction partners of kinases (derived from the STRING database) are more likely to be phosphorylated by the respective kinase, they should therefore contain an amino acid motif conferring kinase specificity. This can then be uncovered by motif enrichment.

The described methods provide predictions that are very valuable but not free from error, for example due the described differences in *in vitro* and *in vivo* kinase specificity or the influence of sub-cellular localization. Thus, the predicted kinase-substrate interactions should be considered hypotheses to be tested experimentally [Linding et al., 2007].

We hereafter present five computational methods to infer kinase activities from phosphoproteomics data, which use either curated or computationally predicted kinase-substrate interactions.

### 3.2 GSEA (Gene Set Enrichment Analysis)

Methodologically, inference of kinase activity from phosphoproteomics data is related to the inference of transcription factor activity based on gene expression data. A plethora of different methods has been developed for the prediction of transcription factor activity, e.g. the classical gene set enrichment analysis [Subramanian et al., 2005] or elaborated machine learning methods [Schacht et al., 2014].

For example, Drake et al. [Drake et al., 2012] analyzed the kinase signaling network in castration-resistant prostate cancer with GSEA. They predicted the kinases responsible for each phosphosite with kinase-substrate interactions from PhosphoSitePlus, kinase recognition motifs from PHOSIDA, and predictions from NetworKIN. Subsequently, they computed the enrichment of each kinases’ targets with the gene set enrichment algorithm after Subramanian et al. [Subramanian et al., 2005], which corresponds to a Kolmogorov–Smirnov-like statistic. The significance of the enrichment score is determined based on permutation tests, whereas the p-value depends on the number of permutations.

Alternatively, the gene set enrichment web-tool Enrichr [Chen et al., 2013; Kuleshov et al., 2016] can also be used for kinases [Lachmann and Ma’ayan, 2009]. The authors compiled kinases-substrate interactions from different databases and extracted additional interactions manually from the literature in order to generate kinase-targets sets. Furthermore, they added protein-protein interactions involving kinases from the Human Protein Reference Database (HPRD) [Keshava Prasad et al., 2009], based on the assumption that those are highly enriched in kinase-substrate interactions. Using this prior knowledge, the enrichment of the targets of a kinase is then computed with Fisher’s exact test as described in [Chen et al., 2013].

### 3.3 KAA (Kinase Activity Analysis)

Another approach to link phosphoproteomics data with the activity of kinases was presented in a publication from Qi et al. [Qi et al., 2014], which they termed kinase activity analysis (KAA).

In this study, the authors collected phosphoproteomics data from adult mouse testis in order to investigate the process of mammalian spermatogenesis. With the software package iGPS [Song et al., 2012] they predicted putative kinase-substrate relationships for the detected phosphosites. The authors hypothesized that the number of links for a given kinase in the predicted kinase-substrate network can serve as proxy for the activity of this kinase in the specific cell type. This activity was then compared to the kinase activity background which was calculated by computing the number of links in the background kinase-substrate network based on the mouse phosphorylation atlas by Huttlin et al. [Huttlin et al., 2010]. Qi and colleagues predicted high activity of PLK kinases in adult mouse testis and could validate this prediction experimentally.

However, there are several limitations of KAA. For once, it is mainly based on computational predictions of kinase substrate relationships, which are known to be susceptible to errors [Newman et al., 2014; Linding et al., 2007]. Additionally, in their method the activity of a kinase is only dependent on the number of detected, putative targets. The abundance of the individual phosphosites or the fold change between conditions is not taken into account.

De Graaf et al. [de Graaf et al., 2014] chose a comparable approach in a study of the phosphoproteome of Jurkat T cells after stimulation with prostaglandin E_2_. However, they did not explicitly calculate kinase activities; rather, they grouped phosphosites in different clusters with distinct temporal profiles and used the NetworKIN algorithm [Linding et al., 2008] to calculate the enrichment of putative targets of a given kinase in a specific cluster. As a result, they associated kinases with temporal activity profiles based on the enrichment in one of the detected clusters.

### 3.4 CLUE (CLUster Evaluation)

A method designed specifically for time-course phosphoproteomics data is the knowledge-based CLUster Evaluation approach, in short CLUE [Yang et al., 2015]. This method is based on the assumption that phosphosites targeted by the same kinase will show similar temporal profiles, which is utilized to guide a clustering algorithm and infer kinases associated to these clusters. As in the study by de Graaf et al. [de Graaf et al., 2014], kinases are not associated with distinct values for activities but rather with temporal activity profiles. The notable distinction of CLUE is that the clustering is found based on the prior-knowledge about kinase-substrate relationships, as outlined below.

Methodologically, CLUE uses the *k*-means clustering algorithm to group the phosphoproteomics data into clusters in which the phosphosites show similar temporal kinetics. The performance of *k*-means clustering is particularly sensitive to the parameter *k*, i.e. the number of clusters. CLUE therefore tests a range of different values for *k* and evaluates them based on the enrichment of kinase-substrate relationships in the identified clusters. The method utilizes the data from the Phospho-SitePlus database in order to derive prior-knowledge about kinase-substrate relationships. With Fisher’s exact test the enrichment of the targets of a given kinase in a specific cluster is tested for significance. The implemented scoring system penalizes distribution of the targets of a single kinase throughout several clusters, as well as the combination of unrelated phosphosites in the same cluster.

CLUE is freely available as R package in the Comprehensive R Archive Network CRAN under https://cran.r-project.org/web/packages/ClueR/index.html.

A limitation of CLUE is represented by the fact that possible ‘noise’ in the prior-knowledge, i.e. incorrect annotations, potentially derived from cell type specific kinase-substrate relationships, can affect the performance of the clustering, although simulations showed reasonable robustness. CLUE is tailored towards time-course phosphoproteomics data and associates kinases with temporal activity profiles. Since the method does not provide singular activity scores for each kinase, it may be only partly applicable to experiments in which the individual responses of kinases to different treatments or conditions are of interest.

### 3.5 KSEA (Kinase Set Enrichment Analysis)

Casado et al. [Casado et al., 2013] presented a method for kinase activity estimation based on kinase substrate sets. Using kinase-substrate relationships derived from the databases PhosphoSitePlus and Phospho.ELM, all phosphosites that are targeted by a given kinase can be grouped together into a substrate set (see Figure 1 for an outline of the work flow). In theory, these phosphosites should show similar values, since they are targeted by the same kinase. However, due to the transient nature of phosphorylation, additional biological variability like cell cycle status, and the possibility that the reported kinase-substrate link may not be present in the specific cell type of interest, phosphoproteomics measurements can be considered inherently noisy. Therefore, Casado and colleagues proposed integrating the information from all phosphosites in the substrate set in order to enhance the signal-to-noise ratio by signal averaging [Wilm and Mann, 1996].

**Figure 1:**
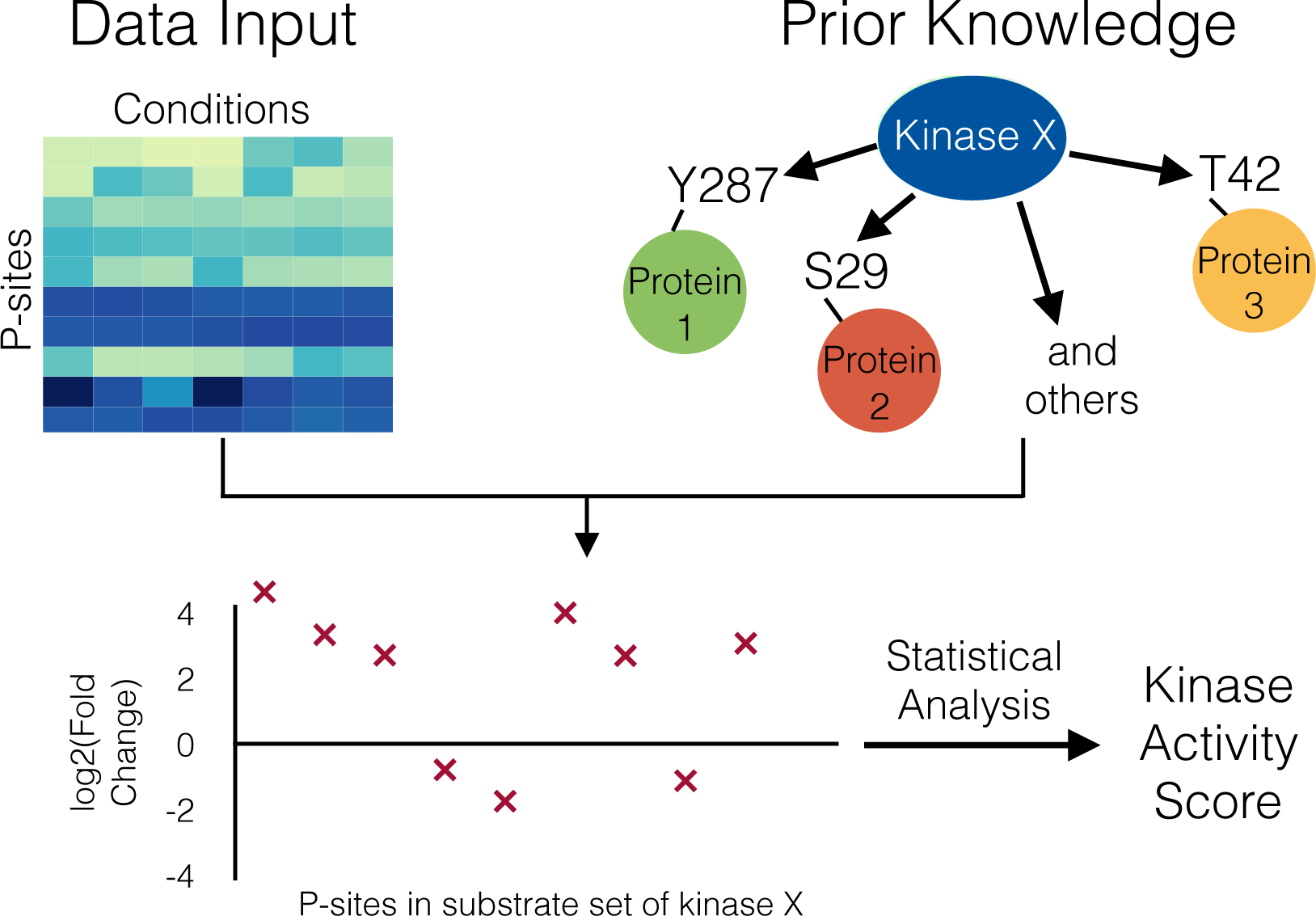
Work-flow of methods to obtain Kinase activity scores such as KSEA. As prior-knowledge, the targets of a given kinase are extracted out of curated databases like PhosphoSitePlus. Together with the data of the detected phosphosites, substrate sets are constructed for each kinase, from which an activity score can be calculated.

For KSEA, log2-transformed fold change data is needed, i.e. the change of abundance in a phosphosite between initial and treated state, initial and later time point, or between two different cell types. Therefore, KSEA activity scores describe the activity of a kinase in one condition relative to another.

The authors suggested three possible metrics (mean score, alternative mean score, and delta score) that can be extracted out of the substrate set and serve as proxy for kinase activity: (i) The main activity score, also used in following publications [Wilkes et al., 2015], is defined as the mean of the log2 fold changes of the phosphosites in the substrate set; (ii) alternatively, only phosphosites with significant fold changes can be considered for the calculation of the mean; and (iii) for the last approach, termed ‘delta count’, the occurrence of significantly up-regulated phosphosites in the substrate set is counted, from which the number of significantly down-regulated sites is subtracted. For each method, the significance of the kinase activity score is tested with an appropriate statistical test. In the publication of Casado et al., all three measures were in good agreement, even if spanning different numerical ranges (see Figure 2). The implementation of these three methods is discussed in detail in the following section.

**Figure 2:**
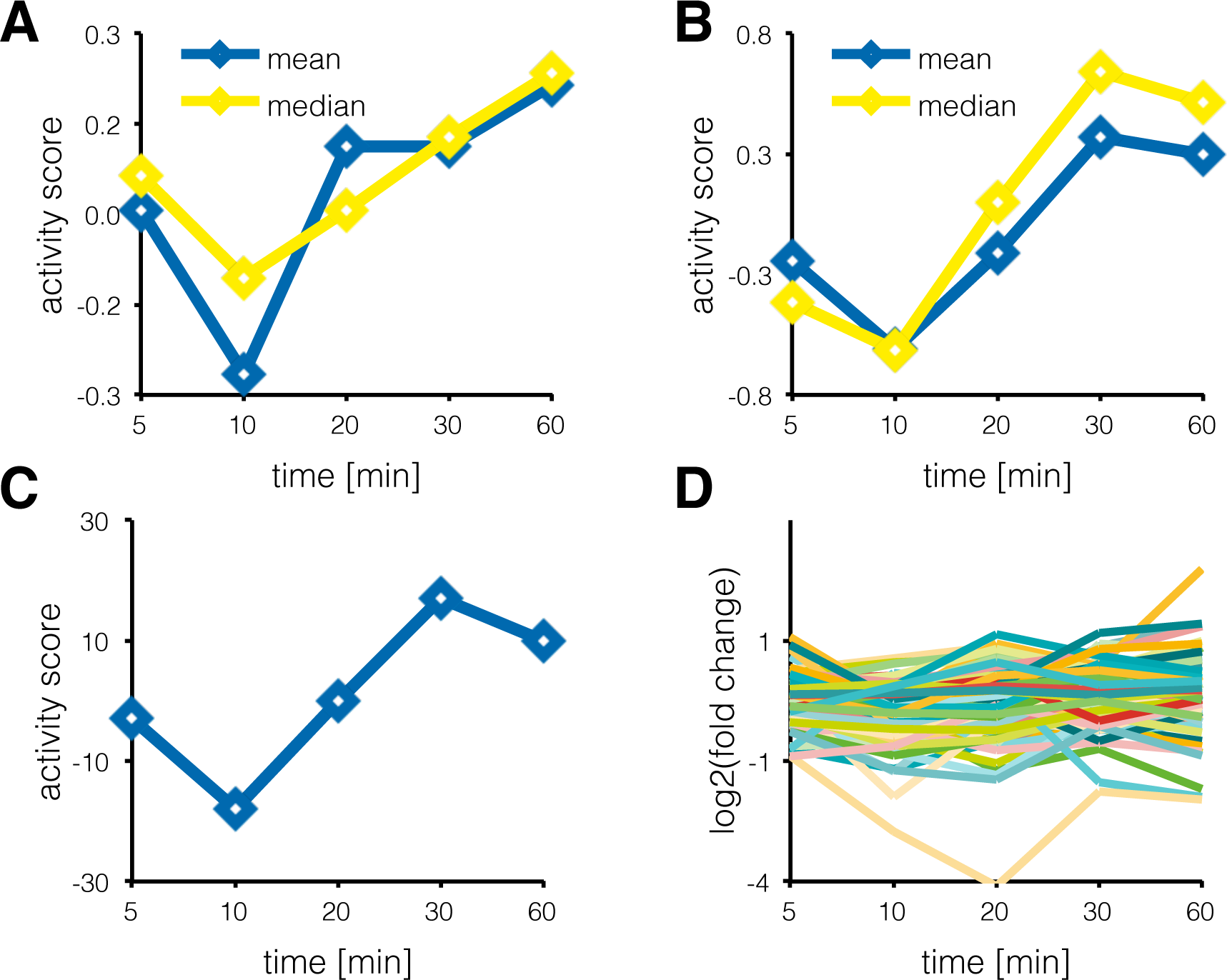
KSEA activity scores for Casein kinase II subunit alpha A: Activity scores for Casein kinase II subunit alpha over all time points of the de Graaf data set [de Graaf et al., 2014], calculated as the mean of all phosphosites in the substrate set. In yellow, the median has been used. B: Activity scores for Casein kinase II subunit alpha over all time points of the de Graaf data set, calculated as the mean of all significantly regulated phosphosites in the substrate set. The median is again shown in yellow. C: Delta score for Casein kinase II subunit alpha over all time points of the de Graaf data set, calculated as number of significantly upregulated phosphosites minus the number of significantly down-regulated phosphosites in the substrate set. D: The log2 fold changes for all time points for all phosphosites in the substrate set of the Casein kinase II subunit alpha.

Like the other methods described in this section, KSEA strongly depends on the prior-knowledge kinase-substrate relationships available in the freely accessible databases. These are far from complete and therefore limit the analytical depth of the kinase activity analysis. Additionally, databases are generally biased towards well-studied kinases or pathways [Horn et al., 2014], so that the sizes of the different substrate sets differ considerably. Casado et al. tried to address these limitations by integrating information about kinase recognition motifs and obtained comparable results.

A detailed protocol on how to use KSEA is provided in the next section.

### 3.6 IKAP (Inference of Kinase Activity from Phosphoproteomics)

Recently, Mischnik and colleagues introduced a machine-learning method to estimate kinase activities and to predict putative kinase-substrate relationships from phosphoproteomics data [Mischnik et al., 2015].

In their model for kinase activity, the effect *e* of a given kinase *j* on a single phosphosite *i* is modeled with

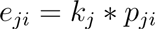

as product of the kinase activity *k* and the affinity *p* of kinase *j* for phosphosite *i*. The abundance *P* of the phosphosite *i* is expressed as mean of all effects acting on it, since several kinases can regulate the same phosphosite:

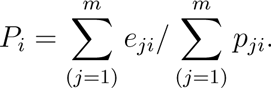

The information about the kinase-substrate relationships is also derived from the PhosphoSitePlus database. Using a non-linear optimization routine, IKAP estimates the described parameters while minimizing a least square cost function between predicted and measured phosphosite abundance throughout time points or conditions. For this optimization, the affinity parameters are estimated globally, while the kinase activities are fitted separately for each time point.

In a second step, putative new kinase-substrate relationships are predicted based on the correlation of a phosphosite with the estimated activity of a kinase throughout time points or conditions. These predictions are then tested by database searches and by comparison to kinase recognition motifs from NetworKIN.

In contrast to KSEA, which computes the kinase activity based on the fold changes of the phosphosites in the respective substrate set, IKAP is built on a heuristic machine learning algorithm and tries to fit globally the described model of kinase activity and affinity to the phosphoproteomics data. Therefore, the output of IKAP is not only a score for the activity of a kinase, but also a value representing the strength of a specific kinase-substrate interaction in the investigated cell type. On the other hand, the amount of parameters that have to be estimated is rather large, so that a fair number of experimental conditions or time points are needed for unique solutions. Mischnik et al. included a function to perform an identifiable analysis of the obtained kinase activities and could show in the case of the two investigated example datasets that the found solutions are indeed unique on the basis of the phosphoproteomics measurements.

The MATLAB code for IKAP can be found online under www.github.com/marcel-mischnik/IKAP, accompanied by an extensive step-by-step documentation, which we recommend as additional reading to the interested reader.

## 4 Protocol for KSEA

In this section, we present a stepwise, guided protocol for the KSEA approach to infer kinase activities from phosphoproteomics data. This protocol (part of the Kinase Activity Toolbox under https://github.com/saezlab/kinact) is accompanied by a freely available script, written in the Python programming language (Python version 2.7.x) that should enable the use of KSEA for any phosphoproteomics dataset. We plan to expand kianct to other methods in the future. We are going to explain the performed computations in detail in the following protocol to facilitate understanding and to enable a potential re-implementation in other programming languages.

As example application, we will use KSEA on the phosphoproteomics dataset from de Graaf et al., 2014 [de Graaf et al., 2014], which was derived from Jurkat T cells stimulated with prostaglandin E_2_ and is available as supplemental information to the article online at http://www.mcponline.org/content/13/9/2426/suppl/DC1.

### 4.1 Quick Start

As a quick start for practiced Python users, we can use the utility functions from kinact to load the example dataset. The data should be organized as Pandas DataFrame containing the log2-transformed fold changes, while the columns represent different conditions or time points and the row individual phosphosites. The p-value of the fold change is optional, but should be organized in the same way as the data.

~~~
import kinact
data_fc, data_p_value = kinact. get_example_data ( )
print data_fc . head ( )
>>>                    5min    10min       20min       30min      60min
>>> ID
>>> A0AVK6_S71   −0.319306   −0.484960   −0.798082   −0.856103   −0.928753
>>> A0FGR8_S743  −0.856661   −0.981951   −1.500412   −1.441868   −0.861470
>>> A0FGR8_S758  −1.445386   −2.397915   −2.692994   −2.794762   −1.553398
>>> A0FGR8_S691   0.271458    0.264596    0.501685    0.461984    0.655501
>>> A0JLT2_S226  −0.080786    1.069710    0.519780    0.520883   −0.296040
~~~

The kinase-substrate relationships have to be loaded as well with the function `get_kinase _targets()`. In this function call, we can specify with the sources-parameter, from which databases we want to integrate the information about kinase-substrate relationships, e.g. PhosphoSitePlus, Phospho.ELM or Signor. The function uses an interface to the `pypath` package, which integrates several resources for curated signaling pathways [Turei et al., 2016] (see Note 1).

~~~
kin_sub_interactions = kinact . get_kinase_targets (sources =[ ‘ all ’ ] )
~~~

An important requirement for the following analysis is, that the structure of the indices of the rows of the data and the prior-knowledge need to be the same (see below for more detail). As an example, KSEA can be performed for the condition of 5 minutes after stimulation in the de Graaf dataset using:

~~~
activities, p_values = kinact . ksea . ksea_mean ( data_fc [ ‘ 5min ’ ],
  kin_sub_interactions, mP=data_fc . values . mean (),
  delta=data_fc . values . std ())
**print** activities . head ()
>>>     AKT1                 0.243170
>>>     AKT2                 0.325643
>>>     ATM                 −0.127511
>>>     ATR                 −0.141812
>>>     AURKA                1.783135
>>>     dtype:  float64
~~~

Besides the data (`data_fc[’5min’]`) and kinase-substrate interactions (`kin_sub_interactions`), the variables `mP` and `delta` are needed to determine the z-score of the enrichment. The z-score builds the basis for the p-value calculation. The p-values for all kinases are corrected for multiple testing with the Benjamini-Hochberg procedure [Benjamini and Hochberg, 2000].

In Figure 2, the different activity scores for the Casein kinase II alpha, which de Graaf et al. had associated with increased activity after prolonged stimulation with prostaglandin E_2_, are shown together with the log2 fold change values of all phosphosites that are known to be targeted by this kinase. For the methods that use the mean, the median as more robust measure can be calculated alternatively. The qualitative changes of the kinase activities (Figure 2A, B, and C) are quite similar regardless of the method, and would not be apparent from looking at any specific substrate phosphosite alone (Figure 2D).

### 4.2 Loading the Data

In the following, we walk the reader step by step through the procedure for KSEA. Firstly, we need to organize the data so that the KSEA functions can interpret it.

In Python, the library Pandas [Mckinney, 2010] provides useful data structures and powerful tools for data analysis. Since the provided script depends on many utilities from this library, we would strongly advice the reader to have a look at the Pandas documentation, although it will not be crucial in order to understand the presented protocol. The library, together with the NumPy [Van Der Walt et al., 2011] package, can be loaded with:

~~~
import pandas as pd
import numpy as np
~~~

The data accompanying the article is provided as Excel spreadsheet and can be imported to python using the pandas `read_excel` function or first be saved as csv-file, using the ‘Save As’ function in Excel in order to use it as described below. For convenience, in the referenced Github repository, the data is already stored as csv-file, so that this step is not necessary. The data can be loaded with the function `read_csv`, which will return a Pandas DataFrame containing the data organized in rows and columns.

~~~
data_raw = pd . read_csv ( ‘FILEPATH’, sep= ’, ’ )
~~~

In the DataFrame object `data_raw`, columns represent the different experimental conditions or additional information and the row’s unique phosphosites. A good way to gain an overview about the data stored in a DataFrame or to keep track of changes are the following functions:

~~~
print data_raw . head ( ) # to show the first five rows of the DataFrame
   or
print data_raw . shape # in order to show the dimensions of the
   DataFrame
~~~

Phosphosites that can be matched to different proteins or several positions within the same protein are excluded from the analysis. In this example, ambiguous matching is indicated by the presence of a semicolon that separates multiple possible identifiers.

~~~
data_reduced = data_raw [∼ data_raw [ ‘ Proteins ’ ] . str . contains ( ’ ; ’ ) ]
~~~

For more convenient data handling, we will index each phosphosite with an unambiguous identifier comprising the UniProt accession number, the type of the modified residue, and the position within the protein. For the example of a phosphorylation of the serine 59 in the Tyrosine-protein kinase Lck, the identifier would be P06239_S59. The identifier can be constructed by concatenating the information that should be provided in the dataset. In the example of de Graaf et al., the UniProt accession number can be found in the columns ‘Proteins’, the modified residue in ‘Amino acid’, and the position in ‘Positions within proteins’. The index is used to access the rows in a DataFrame and will later be needed to construct the kinase-substrate sets. After creation of the identifier, the DataFrame is indexed by calling the function `set_index`.

~~~
data_reduced [ ‘ ID ’ ] = data_reduced [ ’ Proteins ’ ] + ’_ ’ +
   data_reduced [ ‘ Amino acid ’ ] +
   data_reduced [ ‘ Positions within proteins ’ ]
data_indexed = data_reduced . set_index ( data_reduced [ ’ ID ’ ] )
~~~

Mass spectrometry data is usually accompanied by several columns containing additional information about the phosphosite (e.g. the sequence window) or statistics about the database search (for example the posterior error probability), which are not necessarily needed for KSEA. We therefore extract only the columns of interest containing the processed data. In the example dataset, the names of the crucial columns start with `Average`, enabling selection by a simple if statement. Generally, more complex selection of column names can be achieved by regular expressions with the python module `re`.

~~~
data _ intensity = data_indexed [ [x for x in data_indexed if x . startswith
    ( ’ Average ’ ) ] ] # see Note 2
~~~

Now, we can compute the fold change compared to the control, which is the condition of 0 minutes after stimulation. With *log*(*a/b*) = *log*(*a*) − *log*(*b*), we obtain the fold changes by subtracting the column with the control values from the rest using the sub function of Pandas (see Note 3).

~~~
data_fc = data_intensity . sub (data_intensity [ ’ Average Log2 Intensity 0 min ’ ], axis =0)
~~~

Further data cleaning (re-naming columns and removal of the columns for the control time point) results in the final dataset:

~~~
data_fc . columns = [ x . split ( ) [ −1] for x in data_fc ] # Rename columns
data_fc . drop ( ’ 0min ’, axis =1, inplace=True ) # Delete control column
print data_fc . head ( )
>>>                    5min       10min       20min       30min       60min
>>> ID
>>> A0AVK6_S71     −0.319306    −0.484960   −0.798082   −0.856103   −0.928753
>>> A0FGR8_S743    −0.856661    −0.981951   −1.500412   −1.441868   −0.861470
>>> A0FGR8_S758    −1.445386    −2.397915   −2.692994   −2.794762   −1.553398
>>> A0FGR8_S691     0.271458     0.264596    0.501685    0.461984    0.655501
>>> A0JLT2_S226    −0.080786     1.069710    0.519780    0.520883   −0.296040
~~~

If the experiments have been performed with several replicates, statistical analysis enables estimation of the significance of the fold change compared to a control expressed by a p-value. The p-value will be needed to perform KSEA using the ’Delta count’ approach but may be dispensable for the mean methods. The example data set contains a p-value (transformed as negative logarithm with base 10) in selected columns and can be extracted using:

~~~
data_p_value = data_indexed [ [ x for x in data_indexed if x . starts with ( ’ p value ’ ) ] ]
data_p_value = data_p_value . as type ( ’ float ’ ) # see Note 4
~~~

### 4.3 Loading the Kinase-Substrate Interactions

Now, we load the prior knowledge about kinase-substrate relationships. In this example, we use the information provided in the PhosphoSitePlus database (see Note 5), which can be downloaded from the website www.phosphosite.org. The organization of the data from comparable databases, e.g Phospho.ELM, does not differ drastically from the one from PhosphoSitePlus and therefore requires only minor modifications. Using pd.read_csv again, we load the downloaded file with:

~~~
ks _rel = pd . read_csv ( ’ file path ’, sep= ’ \ t ’ ) # see Note 6
~~~

In this file, every row corresponds to an interaction between a kinase and a unique phosphosite. However, it must first be restricted to the organism of interest, e.g. ’human’ or ’mouse’, since the interactions of different organisms are reported together in PhosphoSitePlus.

~~~
ks_rel_human = ks_ rel . loc [ ( ks_ rel [ ’KIN_ORGANISM ’ ] == ’human ’ ) & ( ks _rel [ ’SUB_ORGANISM ’ ] == ’human ’ ) ]
~~~

Next, we again construct unique identifiers for each phosphosite using the information provided in the data set. The modified residue and its position are already combined in the provided data.

~~~
ks_rel_human [ ’ p s i t e ’ ] = ks_rel_human [ ’SUB_ACC_ID ’ ] + ’_ ’ + ks_rel_human [ ’SUB_MOD_RSD’ ]
~~~

Now, we construct an adjacency matrix for the phosphosites and the kinases. In this matrix, an interaction between a kinase and a phosphosite is denoted with a 1, all other fields are filled with a 0. For this, the Pandas function pivot_table can be used:

~~~
ks_rel_human [ ’ value ’ ] = 1 # see Note 7
adj_matrix = pd . pivot _ table ( ks_rel_human, values= ’ value ’, index= ’ psite
 ’, columns= ’GENE ’, fill _ value =0)
~~~

The result is an adjacency matrix of the form *m* × *n* with *m* being the number of phospho-sites and *n* the number of kinases. If a kinase is known to phosphorylate a given phosphosite, the corresponding entry in this matrix will be a 1, otherwise a 0. A 0 does not mean, that there cannot be an interaction between the kinase and the respective phosphosite, but rather that this specific interaction has not been reported in the literature. As sanity check, we can print the number of known kinase-substrate interactions for each kinase saved in the adjacency matrix:

~~~
print adj_matrix .sum (axis =0) . sort_values (ascending=False ) . head ( )
>>> GENE
>>> CDK2                   541
>>> CDK1                   458
>>> PRKACA                 440
>>> CSNK2A1       437
>>> SRC           391
>>> dtype: int 64
~~~

### 4.4 KSEA

In the accompanying toolbox, we provide for each method of KSEA a custom python function that automates the analysis for all kinases in a given condition. Here, however, we demonstrate the principle of KSEA by computing the different activity scores for a single kinase and a single condition. As example, the Cyclin-dependent kinase 1 (CDK1, see Note 8) and the condition of 60 minutes after prostaglandin stimulation shall be used.

~~~
data_condition = data_fc [ ’ 60min ’ ] . copy ( )
p_values = data_p_value [ ’ p value_60vs0min ’ ]
kinase = ’CDK1 ’
~~~

First, we determine the overlap between the known targets of the kinase and the detected phosphosites in this condition, because we need it for every method of KSEA. Now, we benefit from having the same format for the index of the dataset and the adjacency matrix. We can use the Python function intersection to determine the overlap between two sets.

~~~
substrate_set = adj_matrix [kinase].replace ( 0, np . nan ) . dropna ( ) . index #
    see Note 9
detected_p_sites = data_condition . index
intersect = list ( set ( substrate_set ) . intersection ( detected_p_sites) )
print len (intersect)
>>> 114
~~~

#### 4.4.1 KSEA using the ’Mean’ method

For the ’mean’ method, the KSEA score is equal to the mean of the fold changes in the substrate set *mS*.

The significance of the score is tested with a *z*-statistic using

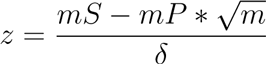

with *m*P as mean of the complete dataset, *m* being the size of the substrate set and the standard deviation of the complete dataset. The ’mean’ method has established itself as the preferred method in the Cutillas lab that developed the KSEA approach.

~~~
mS = data_condition . ix [ intersect ] . mean ( )
mP = data_fc . values . mean ( )
m = len ( intersect ) delta = data_fc . values . std ( )
z_score = (mS −mP) np . sqrt (m) 1/ delta
~~~

The z-score can be converted into a p-value with a function from the SciPy [Jones et al., 2007] library:

~~~
from scipy . stats import norm
p_value_mean = norm . s f ( abs ( z_score ) )
print mS, p_value_mean
>>> −0.441268760191 9.26894825183e−07
~~~

#### 4.4.2 KSEA using the alternative ’Mean’ method

Alternatively, only the phosphosites in the substrate set that change significantly between conditions can be considered when computing the mean of the fold changes in the substrate set. Therefore, we need a cut-off, determining a significant increase or decrease respectively, which can be a user-supplied parameter. Here, we use a standard level to define a significant change with a cut-off of 0.05. The significance of the KSEA score is tested as before with the *z*-statistic.

~~~
cut_off = −np . log10 ( 0.05 )
set_alt = data_condition . ix [ intersect ] . where (
       p_values . ix [ intersect ] > cut_off) . dropna ( )
mS_alt = set_alt . mean ( )
z_score_alt = ( mS_alt −mP) np . sqrt ( len ( set_alt ) ) 1/ delta
p_value_mean_alt = norm . sf ( abs ( z_score_alt ) )
print mS_alt, p_value_mean_alt
>>> −0.680835732551 1.26298232031e−13
~~~

#### 4.4.3 KSEA using the ’Delta count’ method

In the ’Delta count’ method, we count the number of phosphosites in the substrate set that are significantly increased in the condition versus the control and subtract the number of phosphosites that are significantly decreased.

~~~
cut_off = −np . log 10 ( 0.05 )
score_delta = len ( data_condition . ix [ intersect ] . where (
              ( data_condition . ix [ intersect ] > 0) &
              ( p_values . ix [ intersect ] > cut_off ) ) . dropna ( ) ) −
              len ( data_condition . ix [ intersect ] . where (
              ( data_condition . ix [ intersect ] < 0) &
              ( p_values . ix [ intersect ] > cut_off) ) . dropna ( ) ) # see
                 Note 10
~~~

The p-value of the score is calculated with a hypergeometric test, since the number of significantly regulated phosphosites is a discrete variable. To initialize the hypergeometric distribution, we need as variables *M* = the total number of detected phosphosites, *n* = the size of the substrate set, and *N* = the total number of phosphosites that are in an arbitrary substrate set and significantly regulated.

~~~
from scipy . stats import hypergeom
M = len ( data_condition )
n = len ( intersect )
N = len ( np . where ( p_values . ix [ adj_matrix . index . tolist ( ) ] > cut_off) [ 0 ] )
hypergeom_dist = hypergeom (M, n, N)
p_value_delta = hypergeom_dist . pmf ( len ( p_values . ix [ intersect ] . where (
                                  p_values . ix [ intersect ] > cut_off ).
                                     dropna ( ) ) )
print score_delta, p_value_delta
>>> −58 8.42823410966e−119
~~~

## 5 Notes

1. The available sources for kinase-substrate interactions are ARN [Türei et al., 2015], CA1 [Ma’ayan et al., 2005], dbPTM [Huang et al., 2016], DEPOD [Duan et al., 2015], HPRD [Keshava Prasad et al., 2009], MIMP [Wagih et al., 2015], Macrophage [Raza et al., 2010], NRF2ome [Türei et al., 2013], phosphoELM [Dinkel et al., 2011], PhosphoSite [Hornbeck et al., 2015], SPIKE [Paz et al., 2011], SignaLink3 [Fazekas et al., 2013], Signor [Perfetto et al., 2016], and TRIP [Chun et al., 2014].
2. The provided code is equivalent to:

~~~
intensity_columns = [ ]
for x in data_indexed :
        if x . starstwith ( ’ Average ’ ) :
                   intensity_columns . append ( x )
data_intensity = data_indexed [ intensity_columns ]
~~~
3. In our example it is not necessary to transform the data to log2 intensities, since the data is already provided after log2-transformation. But for raw intensity values, the following function from the NumPy module can be used:

~~~
data_log2 = np . log2 ( data_intensity )
~~~
4. Due to a compatibility problem with the output of Excel, Python recognizes the p-values as string variables, not as floating point numbers. Therefore, this line is needed to convert the type of the p-values.
5. The adjacency matrix can also be constructed based on kinase recognition motifs or kinase prediction scores and the amino acid sequence surrounding the phosphosite. To use NetworKIN scores for the creation of the adjacency matrix, kinact will provide dedicated functions. In the presented example, however, we focus on the curated kinase-substrate relationships from PhosphoSitePlus.
6. The file from PhosphoSitePlus is provided as text file in which a tab (`\t`) delimits the individual fields, not a comma. The file contains a disclaimer at the top, which has to be removed first. Alternatively, the option `skiprows` in the function read_csv can be set in order to ignore the disclaimer.
7. This column is needed, so that in the matrix resulting from `pd.pivot_table` the value from this column will be entered.
8. If necessary, mapping between protein names, gene names, and UniProt-Accession numbers can easily be performed with the Python module `bioservices`, to the documentation of which we want to refer the reader [Cokelaer et al., 2013].
9. In this statement, we first select the relevant columns of the kinase from the connectivity matrix (`adj_matrix[kinase]`). In this column, we replace all 0 values with NAs (`replace(0, np.nan)`), which are then deleted with `dropna()`. Therefore, only those interactions remain, for which a 1 had been entered in the matrix. Of these interactions, we extract the index, which is a list of the phosphosites known to be targeted by the kinase of interest.
10. The `where` method will return a copy of the DataFrame, in which for those cases that the condition is not true, a NA is returned. `dropna()` will therefore delete all those occurrences, so that `len(`) will count how often the condition is true.

## 6 Closing Remarks

In summary, the methods described in this review use different approaches to calculate kinase activities or to relate kinases to activity profiles from phosphoproteomic datasets. All of them utilize prior-knowledge about kinase-substrate relationships, either from curated databases or from computational prediction tools. Using these methods, the noisy and complex information from the vast amount of detected phosphorylation sites can be condensed into a much smaller set of kinase activities that is easier to interpret. Modeling of signaling pathways or prediction of drug responses can be performed in a straightforward way with these kinase activities as shown in the study by Casado et al. [Casado et al., 2013].

The power of the described methods strongly depends on the available prior-knowledge about kinase-substrate relationships. As our knowledge increases due to experimental methods like *in vitro* kinase selectivity studies [Imamura et al., 2014] or CEASAR (Connecting Enzymes And Substrates at Amino acid Resolution) [Newman et al., 2013], the utility and applicability of methods for inference of kinase activities will grow as well. Additionally, the computational approaches for prediction of possible kinase-substrate relationships are under on-going development [Wagih et al., 2016; Creixell et al., 2015a], increasing the reliability of the *in silico* predictions.

Phosphoproteomic data is not only valuable for the analysis of kinase activities: for example, PTMfunc is a computational resource that predicts the functional impact of post-translational modifications based on structural and domain information [Beltrao et al., 2012], and PHONEMeS Wilkes et al. [2015]; Terfve et al. [2015] combines, similar to kinase-activity methods, phosphoproteomics data with prior-knowledge kinase-substrate relationships. However, instead of scoring kinases, PHONEMeS derives logic models for signaling pathways at the phosphosite level.

For the analysis of deregulated signaling in cancer, mutations in key signaling molecules can be of crucial importance. Recently, Creixell and colleagues presented a systematic classification of genomic variants that can perturb signaling, either by rewiring of the signaling network or by destruction of phosphorylation sites [Creixell et al., 2015b]. Another approach was introduced in the last update of the PhosphoSitePlus database, in which the authors reported with PTMVar [Hornbeck et al., 2015] the addition of a dataset that can map missense mutation onto the post-translational modifications. With these tools, the challenging task of creating an intersection between genomic variations and signaling processes may be addressed.

It remains to be seen how the different scoring metrics for kinase activity relate to each other, as they utilize different approaches to extract a kinase activity score out of the data. IKAP is based on a non-linear optimization for the model of kinase-dependent phosphorylation, KSEA on statistical analysis of the values in the substrate set of a kinase, and CLUE on the k-means clustering algorithm together with Fisher’s exact test for enrichment. To assess the different methods, they have to be benchmarked against ’gold standard’ datasets for which the activity status of kinases is known. Such comprehensive comparison of applicability, performance and drawbacks for the different methods would be very valuable for the most effective use of phosphoproteomic data to infer kinase activities, from which to derive insights into molecular cancer biology and many other processes controlled by signal transduction.

## A Acknowledgements

Thanks to Emanuel Gonçalves, Aurélien Dugourd, and Claudia Hernández-Armenta for helpful comments on the manuscript.

For help with the code, thanks to Emanuel Gonçalves.

